# RG motifs promote piRNA-mediated gene silencing in *C. elegans*

**DOI:** 10.1101/2025.05.12.653514

**Authors:** Dylan C. Wallis, Carolyn M. Phillips

## Abstract

Argonaute proteins are essential players in RNA silencing pathways, and their N-terminal extensions, particularly in the PIWI clade, often harbor conserved sequences like RG motifs. Despite their prevalence in Argonaute proteins, the role of these motifs remains poorly understood. In this study, we focus on the RG motifs within the N-terminal region of *Caenorhabditis elegans* PRG-1, a PIWI clade Argonaute. Using sequence alignment across Caenorhabditis species, we identify three conserved RG motifs that are methylated, as confirmed by mass spectrometry. The region surrounding these motifs is intrinsically disordered, as predicted by disorder algorithms and structural modeling. While the RG motifs are critical for fertility, germline morphology, and piRNA silencing, they are not required for PRG-1 expression, localization, or piRNA loading. Notably, mutation of the RG motifs results in defects in downstream small RNA production, specifically the depletion of mutator class siRNAs, without affecting piRNA biogenesis. These findings suggest that the RG motifs of PRG-1 play a crucial role in linking piRNA-mediated silencing to siRNA production, and that their function is critical for fertility and germline maintenance in *C. elegans*. Despite these defects, the phenotypic severity in the RG mutant is milder than in a PRG-1 null mutant, highlighting the complexity of PRG-1 function and its post-translational modifications.

## Introduction

Piwi-interacting RNAs (piRNAs) are a class of small non-coding RNAs that associate with Piwi-clade Argonaute proteins, playing a crucial role in germ cell function across metazoans (Aravin et al., 2006; Girard et al., 2006; Grivna et al., 2006; Houwing et al., 2007; Lau et al., 2006; Vagin et al., 2006). Like other small RNA classes, piRNAs target transcripts with full or partial complementarity and mediate gene silencing through mechanisms such as target transcript cleavage, or the recruitment of RNA-dependent RNA polymerases and chromatin-modifying enzymes (Almeida et al., 2019; Czech and Hannon, 2016; Wang et al., 2023). One of the primary roles of piRNAs is the silencing of transposons. By targeting transposon-derived RNAs for degradation or transcriptional silencing, piRNAs help maintain genome stability and integrity (Batista et al., 2008; Carmell et al., 2007; Cox et al., 1998; Deng and Lin, 2002; Lin and Spradling, 1997; Wang and Reinke, 2008). However, emerging evidence also suggests that piRNAs regulate endogenous transcripts, contributing to gene expression control, heterochromatin formation, and the maintenance of fertility (Bagijn et al., 2012; Barckmann et al., 2015; Goh et al., 2015; Gou et al., 2014; Lee et al., 2012; Shirayama et al., 2012).

In *Caenorhabditis elegans*, piRNAs are referred to as 21U-RNAs, named for their characteristic length of 21 nucleotides (nt) and strong 5’ uridine preference (Ruby et al., 2006). A single functional Piwi homolog, Piwi Related Gene-1 (PRG-1), binds to thousands of unique *C. elegans* piRNAs (Batista et al., 2008; Das et al., 2008; Wang and Reinke, 2008). The precursors of mature piRNAs are transcribed by RNA Polymerase II as individual units, with the majority located in two large clusters on Chromosome IV (Gu et al., 2012; Ruby et al., 2006). These "Type I" piRNAs contain an upstream 8-nt Ruby motif and require a complex of proteins, including PRDE-1 and SNPC-4, for their transcription (Kasper et al., 2014; Ruby et al., 2006; Weick et al., 2014). "Type II" piRNAs, found outside of the Chromosome IV clusters, are transcribed from promoters throughout the genome (Gu et al., 2012). After transcription, single-stranded piRNA precursors undergo a series of nucleolytic processing steps before being bound by PRG-1 in the perinuclear germ granules (Batista et al., 2008; Podvalnaya et al., 2023; Tang et al., 2016; Wang and Reinke, 2008). Within these granules, piRNAs are thought to target nascent transcripts as they exit the nuclear pore, triggering the production of additional small RNAs and leading to transcriptional and translational gene silencing.

In most organisms, piRNAs are amplified through the ping-pong cycle, but this mechanism is absent in *C. elegans* (Aravin et al., 2008, 2007; Brennecke et al., 2007; Gunawardane et al., 2007). Instead, efficient piRNA-mediated silencing in *C. elegans* relies on amplification of the signal via the mutator complex. In this process, RNA-dependent RNA polymerases (RdRPs) are recruited to target transcripts, generating anti-sense small interfering RNAs (siRNAs) that are approximately 22 nucleotides long and exhibit a 5’ guanosine bias (22G-RNAs). These 22G-RNAs are then loaded onto worm-specific Argonaute proteins (WAGOs), leading to a robust transcriptional and post-transcriptional silencing response (Yigit et al., 2006; Phillips et al., 2012; Gu et al., 2009; Vasale et al., 2010; Gent et al., 2010; Bagijn et al., 2012). Despite a detailed understanding of the mutator complex, the precise connection between the piRNA pathway and the mutator pathway remains poorly understood.

Argonaute and Piwi proteins can be modified to modulate their biochemical activities. For example, phosphorylation of the *C. elegans* microRNA (miRNA) Argonaute, ALG-1, is critical for miRNA-mediated gene silencing (Quévillon Huberdeau et al., 2022, 2017). Similarly, human AGO2 is highly phosphorylated and at least one phosphorylation site is critical for miRNA binding, while others are critical for Argonaute-mRNA target interactions (Quévillon Huberdeau et al., 2017; Rüdel et al., 2011). Further, many of the miRNA Argonaute phosphorylation sites are conserved between species, including human, mouse, rat, zebrafish, and *C. elegans* (Quévillon Huberdeau et al., 2017). Other modification such as ubiquitylation, Poly(ADP-ribose) modification, and hydroxylation have also been shown to play roles in homeostatic control of small RNA pathways, regulation of miRNAs during stress, subcellular localization of Argonaute proteins, and selection of alternate mRNA silencing modes (Jee and Lai, 2014; Leung et al., 2011; Qi et al., 2008; Quévillon Huberdeau et al., 2022, 2017; Smibert et al., 2013; Wu et al., 2011).

RG and RGG motifs are repetitive amino acid sequences rich in arginine (R) and glycine (G) residues, found in proteins that perform a variety of critical functions, including nucleic acid binding, protein-protein interactions, and protein localization (Thandapani et al., 2013). These motifs can also serve as platforms for arginine methylation, a modification where either one or two methyl groups are added to the terminal amines of the arginine side chain (Bedford and Clarke, 2009). This process is mediated by protein arginine methyltransferases (PRMTs), which are classified into subgroups based on their catalytic activity: type I enzymes catalyze asymmetric dimethylation, type II enzymes catalyze symmetric dimethylation, and type III enzymes catalyze monomethylation (Thandapani et al., 2013). Piwi Argonaute proteins from mice, frogs, and flies are dimethylated by the PRMT protein PRMT5 (Kirino et al., 2009; Reuter et al., 2009; Vagin et al., 2009), which often enables the Piwi protein to interact with members of the Tudor domain protein family (H. Liu et al., 2010; K. Liu et al., 2010; Nishida et al., 2009; Reuter et al., 2009; Vagin et al., 2009; Webster et al., 2015). Tudor domains are protein-protein interaction modules that recognize methylated arginines, allowing for protein-protein interactions in a methylation-specific manner. In both mice and *Drosophila*, Tudor domain proteins play crucial roles in piRNA accumulation, mRNA target regulation, and PIWI protein localization (Anand and Kai, 2011; Andress et al., 2016; Handler et al., 2011; Huang et al., 2021; Nishida et al., 2009; Reuter et al., 2009; Vagin et al., 2009; Zhang et al., 2011).

In *C. elegans* there are six ORFs with homology to mammalian PRMTs with roles that include lifespan regulation, suppression of DNA damage-induced apoptosis, and modulation of chemosensory and locomotory behaviors (Likhite et al., 2015; Takahashi et al., 2011; Wang and Li, 2012; Yang et al., 2009). We have sought to understand the role of arginine methylation and RG motifs in *C. elegans* RNA silencing pathways, and therefore focused our efforts on identifying *C. elegans* Argonaute proteins with RG motifs and investigating their function. In previous work, we found that the long isoform of the germline Argonaute CSR-1 (CSR-1A) contains methylated RG motifs, and that these motifs are required for small RNA binding specificity (Nguyen and Phillips, 2021). While the precise mechanism by which methylated RG motifs promote CSR-1A small RNA specificity has not yet been determined, it is worth noting that numerous Tudor domain-containing proteins have been identified to play a role in small RNA silencing pathways including EKL-1, ERI-5, RSD-6, TOFU-6, and SIMR-1 (Cordeiro Rodrigues et al., 2019; Duchaine et al., 2006; Goh et al., 2014; Gu et al., 2009; Manage et al., 2020; Thivierge et al., 2011; Tijsterman et al., 2004).

In this study, we aim to explore the functional significance of the conserved RG motifs within the N-terminal extension of PRG-1. Through a combination of sequence analysis, structural predictions, and functional assays, we investigate whether these motifs are essential for the proper function of PRG-1 in the piRNA pathway. Specifically, we focus on the role of RG motifs in the regulation of PRG-1 localization, stability, and its involvement in piRNA-mediated gene silencing. Furthermore, we assess the impact of RG motif mutations on germline morphology, fertility, and small RNA production. Our findings provide novel insights into the molecular mechanisms by which PRG-1 contributes to RNA silencing and fertility, highlighting the importance of post-translational modifications in Argonaute function.

## Results

### PRG-1 contains methylated RG motifs

The four structured domains of Argonaute proteins—N, PAZ, MID, and PIWI—are highly conserved (Sheu-Gruttadauria and MacRae, 2017). However, many Argonaute proteins, including those in the PIWI clade, contain N-terminal extensions of varying lengths and sequence content whose function is less well understood (Huang et al., 2021; Martín-Merchán et al., 2024, 2023). To identify functional sequences in the N-terminal extension of *C. elegans* PRG-1, we first generated a DNA sequence alignment of the first 50 amino acids of *C. elegans* PRG-1 with the PRG-1 orthologs in eight additional *Caenorhabditis* species (Madeira et al., 2024). Despite a lack of annotated domains, we found this region to be highly conserved (Fig. 1A). This region also contains three tandem RG motifs, which are present in eight of the nine *Caenorhabditis* PRG-1 genes. The exception is *C. japonica*, which has only two tandem RG motifs (Fig. 1A). To further investigate the structure of this N-terminal region of PRG-1, we sought to determine whether it contains regions of intrinsic disorder, suggesting that it may not be part of a folded domain. Using the disorder prediction algorithm IUPred2 (Dosztányi et al., 2005; Mészáros et al., 2018), we found that amino acids 1-25 are highly disordered and likely constitute an intrinsically disordered region (IDR) (Fig. 1B). This result is further supported by the predicted protein structure, as determined by Alphafold2 (Jumper et al., 2021; Varadi et al., 2024), which finds that the first ∼30 amino acids of *C. elegans* PRG-1 are unstructured (Fig. S1A).

**Figure 1.**
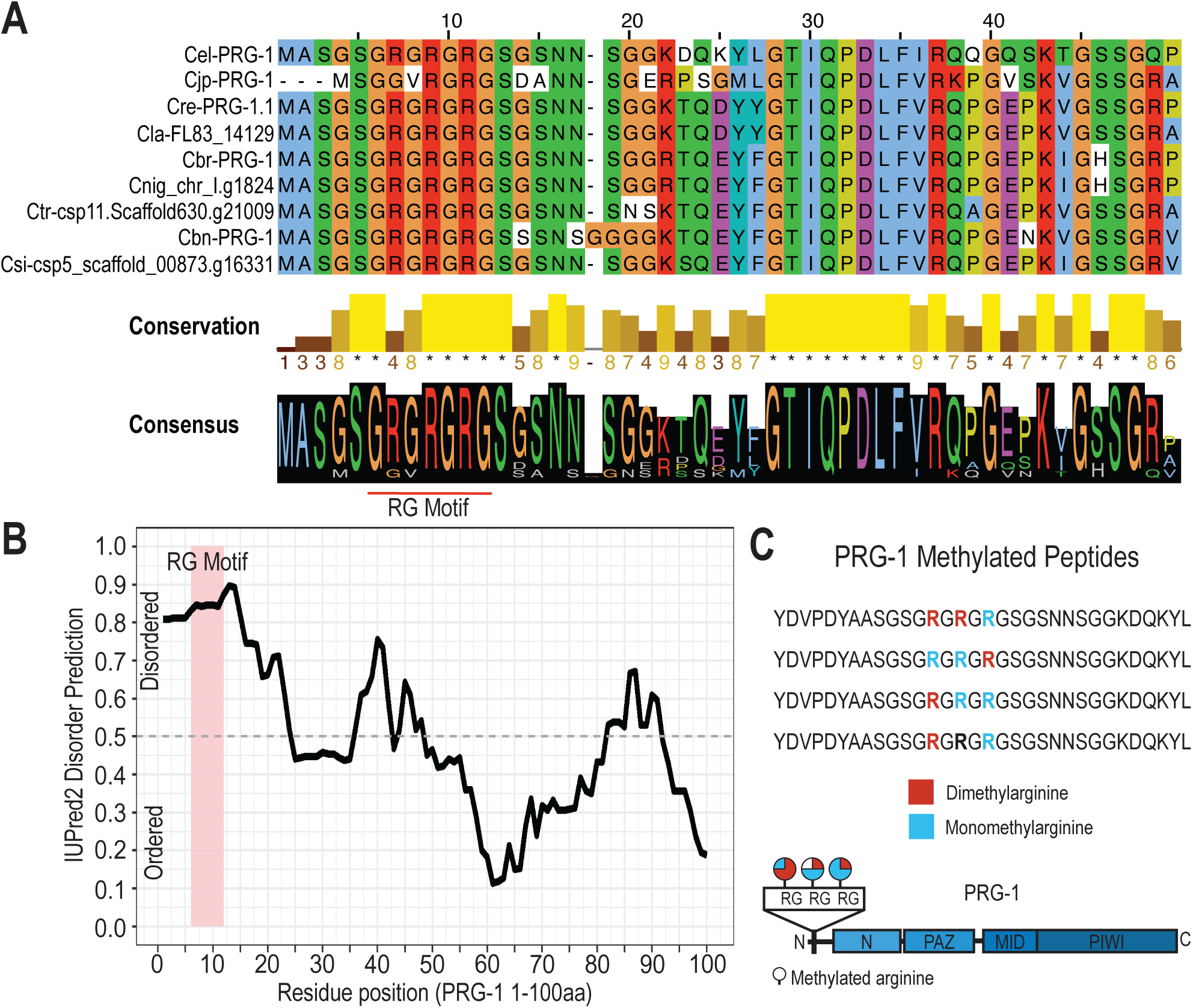
PRG-1 N-terminal extension is unstructured and contains conserved RG motifs. **A.** Clustal Omega Alignment of amino acids 1-50 of *C. elegans* PRG-1 protein sequence and orthologs from the following *Caenorhabditis* species: *japonica*, *remanei*, *latens*, *briggsae*, *nigoni*, *tropicalis*, *brenneri*, and *sinica*. Asterisk (*) in conservation score indicates complete conservation and numbers indicate degree of conservation. Consensus sequence is shown below, where height of amino acid abbreviation is proportional to its abundance at the given position and RG motifs are underlined. **B.** Disorder tendency of the N-terminal region (1-100) of PRG-1 was predicted using IUPred2. **C**. 4 methylated peptides were identified following PRG-1 IP/mass spectrometry (top). Arginines were either dimethylated (red), monomethylated (blue) or unmethylated (black). Diagram below shows schematic of the 3 methylated RG motifs and the frequency each was found to be monomethylated or dimethylated.

Previous studies in *Drosophila*, *Xenopus*, and mouse have found that RG motifs in Piwi orthologs are often dimethylated (Kirino et al., 2009; Reuter et al., 2009; Vagin et al., 2009). In an effort to determine if PRG-1 RG motifs are being post-translationally modified, we immunoprecipitated PRG-1 and subjected it to mass spectrometry analysis. We identified four methylated peptides in this experiment, with all four encompassing the 3 RG motifs located in the N-terminal region of PRG-1 (Fig. 1C). We detected various combinations of mono- and dimethylarginines in these peptides, with each having at least two of the three arginines methylated. Symmetric and asymmetric methylation are mediated by different Protein Arginine Methyltransferase (PRMT) enzymes, are preferentially bound by different interacting proteins, and can have opposite biological effects (Chen et al., 2011; Hartel et al., 2020; H. Liu et al., 2010; K. Liu et al., 2010; Sikorsky et al., 2012; Sun et al., 2011; Thandapani et al., 2013). For example, asymmetric and symmetric dimethylation of histone H3 arginine 2 promote heterochromatin and euchromatin formation, respectively (Hartel et al., 2020; Kirmizis et al., 2007; Migliori et al., 2012). With mass spectrometry, we are unable to distinguish between these isomeric protein modifications. Previous studies have used symmetric and asymmetric methylation-specific antibodies to determine methylation state of target proteins (Kirino et al., 2010, 2009), however these antibodies did not work in our hands. Thus, we could not distinguish between these two variants of the dimethylation leaving this question as a potential avenue for further study. Nonetheless, our data indicate that the N-terminal region of PRG-1 is conserved across the *Caenorhabditis* genus and that multiple methylated RG motifs are present in this unstructured region.

### RG motifs are not required for localization or expression of PRG-1

In order to understand the role of the PRG-1 RG motifs, we generated mutations in all three RG motifs in the first exon of PRG-1, using CRISPR to mutate the three arginine residues to alanine. This mutation will hereafter be referred to as *prg-1(3xAG)*. Next, to determine whether RG motifs are important for germ granule localization, we examined the expression of endogenously-tagged PRG-1 and PRG-1[3xAG]. We found that both PRG-1 and PRG-1[3xAG] localize to perinuclear germ granules in adult animals, with the PRG-1[3xAG] mutant showing no apparent change in morphology or expression level (Fig. 2A). Recently published work has found that PRG-1 accumulates in both P and Z compartments of the germ granule, with an enrichment for the Z granule (Huang et al., 2025). To determine if the RG motifs are required for proper distribution of PRG-1 amongst the different germ granule compartments, we compared PRG-1 localization to that of PGL-1, to mark the P compartment, and ZNFX-1, to mark the Z compartment. We found that both PRG-1 and PRG-1[3xAG] still associate with both P and Z compartments, showing a higher degree of association with the Z compartment (Fig. 2A-B). We next asked if the RG motifs are required for proper expression or stability of PRG-1. By western blot, we determined that PRG-1[3xAG] was expressed at comparable levels to wild-type (Fig. 2C). Together, these experiments demonstrate that the RG motifs in the PRG-1 protein are not required for either expression, stability, or localization of PRG-1.

**Figure 2.**
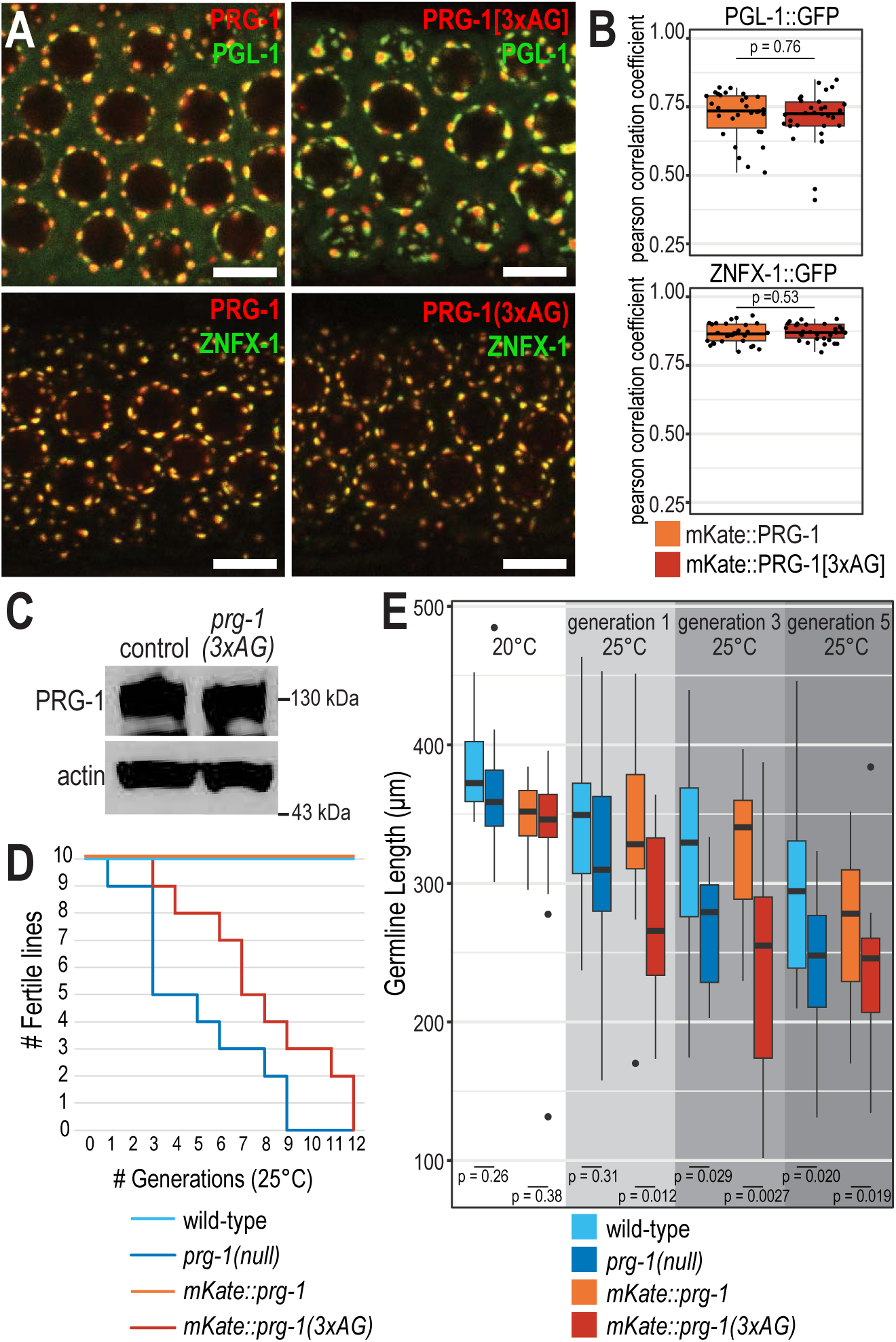
PRG-1 RG motifs are required for transgenerational fertility and germline morphology. **A.** Live imaging of mKate2::PRG-1 or mKate2::PRG-1[3xAG] and GFP::PGL-1 or GFP::ZNFX-1 showing that loss of RG motifs does not affect PRG-1 localization. Scale bars, 5µm. Box plot of Pearson’s R value quantifications between mKate2::PRG-1 (orange) or mKate2::PRG-1[3xAG] (red) and PGL::GFP (top) or GFP::ZNFX-1 (bottom). At least 30 total nuclei from at least 10 individual gonads were used for quantification. Each dot represents an individual quantification, and all data points are shown. Box indicates the first and third quartiles, and whiskers represent the most extreme data points within 1.5 times the interquartile range. Two-tail *t*-tests were performed to determine statistical significance. See materials and methods for a detailed description of quantification methods. **C.** Western blot detecting PRG-1 at gravid adult stage in mKate2::3xMyc::PRG-1 (control) and mKate2::3xMyc::PRG-1[3xAG] strain using α-Myc antibody to detect PRG-1. Actin is shown as a loading control. **D.** Generational fertility of wild-type, *prg-1(null)*, *mKate2::prg-1*, and *mKate2::prg-1(3xAG)* strains. 10 lines for each strain were scored for fertility across 12 generations. The number of lines that were fertile at each generation are shown. **E.** Germline length as measured from distal tip to gonad bend, at 20°C and 1, 2, or 3 generations at 25°C in wild-type, *prg-1(null)*, *mKate2::prg-1*, and *mKate2::prg-1(3xAG)* strains. Box indicates the first and third quartiles, and whiskers represent the most extreme data points within 1.5 times the interquartile range. Outliers are shown as individual points. Two-tail *t*-tests were performed to determine statistical significance.

### PRG-1 RG motifs are required for transgenerational germline morphology and fertility

Proper functioning of the piRNA pathway is critical for maintaining fertility across generations. Depletion of *prg-1*, the sole *C. elegans* PIWI clade Argonaute, results in a progressive sterility (known as a <u>M</u>o<u>rt</u>al Germline or Mrt) that is further exacerbated by elevated temperatures (Simon et al 2014). To investigate whether the PRG-1 RG motifs are required for maintenance of fertility at elevated temperature, we raised ten independent lines of wild-type, *prg-1(null)*, *mKate2::prg-1*, and *mKate2::prg-1(3xAG)* animals at 25°C. As previously shown, *prg-1(null)* animals become progressively sterile (Simon et al., 2014), with all lines sterile by generation 9 (Fig. 2D). Similarly, *prg-1(3xAG)* animals exhibited a progressive sterility phenotype, with all lines sterile by generation 12 (Fig. 2D). These data demonstrate that PRG-1 RG motifs are critical for transgenerational fertility.

It has been shown that the transgenerational fertility defect observed in *prg-1(null)* animals is often associated with shortened or atrophied germlines (Heestand et al., 2018). To further investigate whether the transgenerational fertility defect we observe in *prg-1(3xAG)* mutant animals is similarly associated with germline atrophy, we calculated the mean length of gonad arm, measured from the mitotic tip to the bend of the gonad, for wild-type, *prg-1(null)*, *mKate2::prg-1*, and *mKate2*::*prg-1(3xAG)* animals raised at 20°C, and for 1, 3, or 5 generations at 25°C. The fluorescently-tagged germ granule protein, PGL-1, was included in all strains as a marker for germ cells to simplify quantification. At 20°C, there was no significant difference in germline length for *prg-1(null)* and *mKate2*::*prg-1(3xAG)*, compared to their respective wild-type and *mKate2::prg-1* controls. Following the shift to 25°C, all strains showed some reduction in germline length that became more severe with increasing generations at 25°C, but the *prg-1(null)* and *prg-1(3xAG)* exhibited more significant atrophy than the controls (Fig. 2E). This germline atrophy coincides with the increase in sterility that we observed at generations 3 and 5 for *prg-1(null)* and *prg-1(3xAG)* (Fig. 3D-E). Together, these data indicate that despite the proper expression and localization of PRG-1[3xAG], the RG motifs are required for germ cell maintenance at elevated temperature.

**Figure 3.**
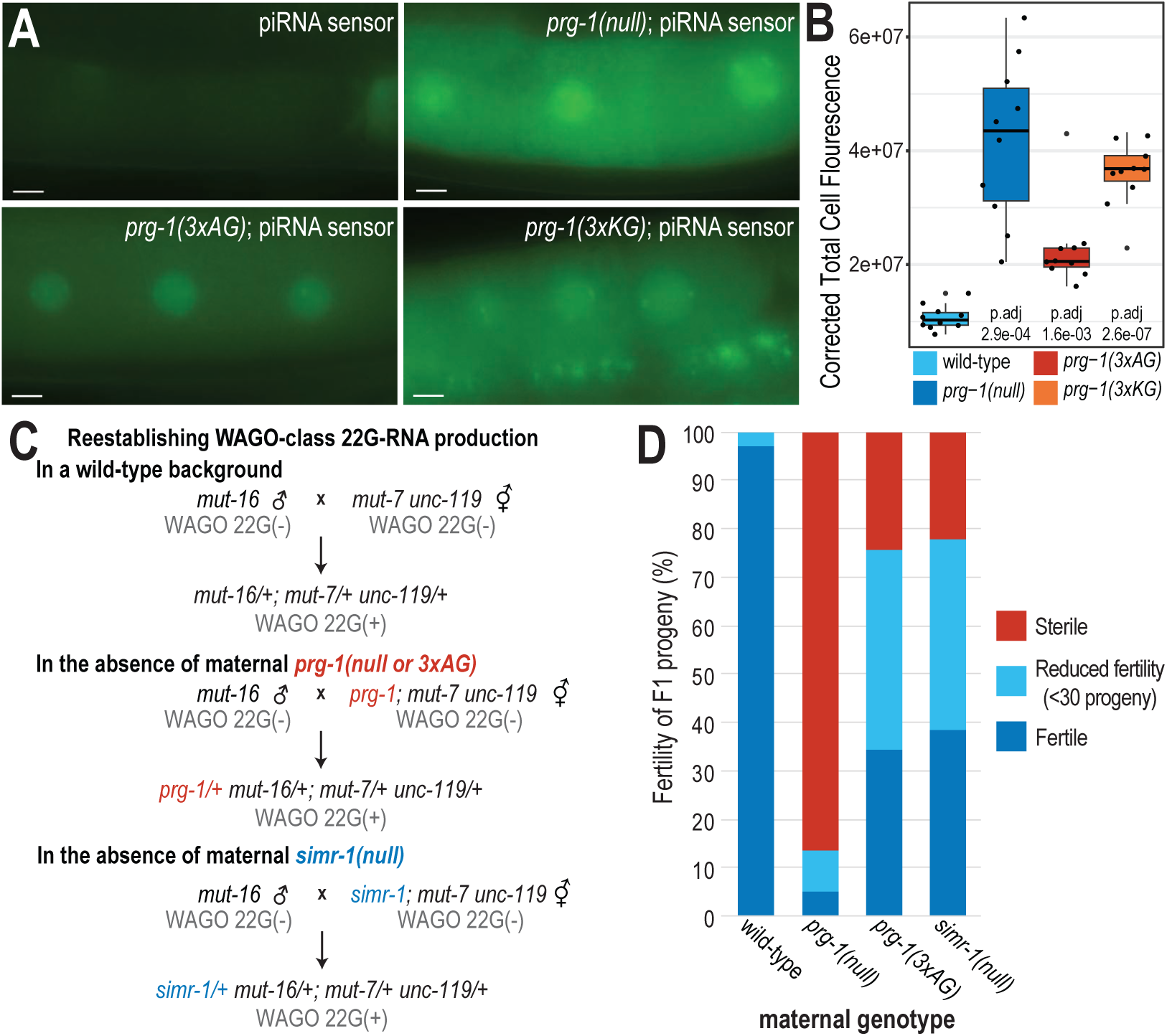
Loss of PRG-1 RG motifs leads to defects associated with piRNA-mediated gene silencing. **A.** Live imaging of the piRNA sensor (*oma-1::gfp; cdk-1::gfp; 21ur-gfp*) in wild-type, *prg-1(null)*, *prg-1(3xAG)*, or *prg-1(3xKG)* mutant backgrounds. Scale bars, 25µm. **B.** Quantification of the piRNA sensor in wild-type, *prg-1(null)*, *prg-1(3xAG)*, or *prg-1(3xKG)* mutant backgrounds. 10 individual -1 oocyte nuclei, one each from 10 individuals were used in quantification. Two-tail *t*-tests were performed to determine statistical significance and *p* values were adjusted for multiple comparisons. Adjusted *p* values for wild-type compared to *prg-1(null)*, *prg-1(3xAG)*, and *prg-1(3xKG)* are shown. **C.** Schematic describing the mating-based approach to reestablish WAGO-class 22G-RNA production in the presence or absence of *prg-1* and *simr-1*. **D.** Bar graph showing the percentage of fertile, sub-fertile (<30 progeny), or sterile F1 animals from the WAGO-class 22G-RNA reestablishment crosses shown in C.

### PRG-1 RG motif mutants exhibit defects in the piRNA pathway

A strain carrying *cdk-1::gfp*, *oma-1::gfp,* and an artificial piRNA targeting GFP can be used to identify genetic perturbations that affect piRNA silencing (Brown et al., 2023). These two reporters show distinct capacities to be targeted for piRNA-mediated silencing, and it has been hypothesized that, when together in the same strain, their silenced state is not robust due to competition between silencing and anti-silencing signals (Brown et al., 2023). Thus, this “piRNA sensor strain” is particularly sensitive to loss of *prg-1*. To investigate whether the RG motifs of PRG-1 are required for piRNA-mediated gene silencing, we introduced the *prg-1(3xAG)* mutation into the piRNA sensor strain using CRISPR. To more directly address whether the methylation of the RG motifs is important for piRNA-mediated gene silencing, we also generated the arginine to lysine mutations in the PRG-1 RG motifs (*prg-1(3xKG)*), which are expected to abolish RG motif methylation while maintaining the positive charge of the motifs. As a control, we also introduced a *prg-1* frameshift mutation (*prg-1(null)*), to completely abolish piRNA activity. We found that *prg-1(null)*, *prg-1(3xAG),* and *prg-1(KG)* mutations all caused significant desilencing of the reporter compared to the unedited strain (Fig. 3A-B). These results indicate that the PRG-1 RG motifs, and specifically methylation of those RG motifs, are required for piRNA-mediated gene silencing.

Another assay that can be used to detect piRNA pathway defects takes advantage of the fact that piRNAs are required to maintain fertility following the loss and subsequent reestablishment of the downstream, WAGO-class 22G-RNA pathway (de Albuquerque et al., 2015; Phillips et al., 2015). In previous work, we have found that if either *prg-1* or the Tudor domain-containing gene, *simr-1,* are disrupted during the reestablishment WAGO-class 22G-RNA production, sterility ensues, resulting from the misrouting of essential genes into the WAGO-dependent silencing pathway (Manage et al., 2020; Phillips et al., 2015). These data indicates that piRNAs are critical to prevent aberrant silencing and misrouting of essential genes. To determine whether the RG motifs of PRG-1 are required for fertility following reestablishment of WAGO-class 22G-RNA production, we set up a reestablishment assay with two mutations in the mutator pathway, *mut-7(-)* and *mut-16(-)*, which are each required for WAGO-class 22G-RNA production. When we cross these two strains, the mutations complement one another, such that the F1 progeny are competent to produce WAGO-class 22G-RNAs but would not inherit WAGO-class 22G-RNAs from their homozygous mutant parents (Fig. 3C). In an otherwise wild-type background, we found that none of the F1 progeny of the reestablishment cross were sterile, although 3% exhibited extremely reduced fertility (a brood size of less than 30). As previously shown, when *prg-1(null)* in is present in maternal parent during the reestablishment cross, the percentage of sterile F1 progeny increased drastically (86% sterile), with an additional 8% exhibiting extremely reduced fertility. Similarly, when the *prg-1(3xAG)* mutant, lacking the PRG-1 RG motifs, was present in the maternal parent during the reestablishment cross 25% of F1 progeny were sterile and 41% exhibited extremely reduced fertility. The fertility defects observed during the reestablishment cross for *prg-1(3xAG)* are that of *simr-1,* where 22% of F1 progeny were sterile and 39% exhibited extremely reduced fertility (Fig. 3D) (Manage et al., 2020). These results indicate that the RG motifs of PRG-1 are required during reestablishment of the WAGO-class 22G-RNA pathway to promote fertility. Further, in conjunction with the requirement of the RG motifs for maintaining silencing of the piRNA sensor, these data indicate that the RG motifs of PRG-1 are critical for proper function of PRG-1 in the piRNA pathway.

### PRG-1 RG motifs are not required for piRNA production or loading

Loss of PRG-1 leads to a dramatic reduction of piRNAs in the germline (Batista et al., 2008; Das et al., 2008; Wang and Reinke, 2008). We next sought to investigate the importance of the RG motifs of PRG-1 for piRNA production and loading. To address the role of the RG motifs of PRG-1 in piRNA production and stability, we sequenced small RNA populations from wild-type, *prg-1(null)*, and *prg-1(3xAG)* mutant animals at 20°C and 25°C. First, we first quantified the total piRNA levels in reads per million (RPM) for each library. As expected, piRNAs were drastically reduced in *prg-1(null)* animals at both temperatures. In contrast, the total piRNA levels in *prg-1(3xAG)* mutants were indistinguishable from wild-type animals (Fig. 4A). Next, we assessed the number of reads mapping to each annotated piRNA locus. Similarly, *prg-1(3xAG)* mutants showed no reduction in piRNA abundance, while *prg-1(null)* animals exhibited a severe reduction in piRNA levels (Fig. 4B). Together, these data demonstrate that the PRG-1 RG motifs are not required for piRNA production.

**Figure 4.**
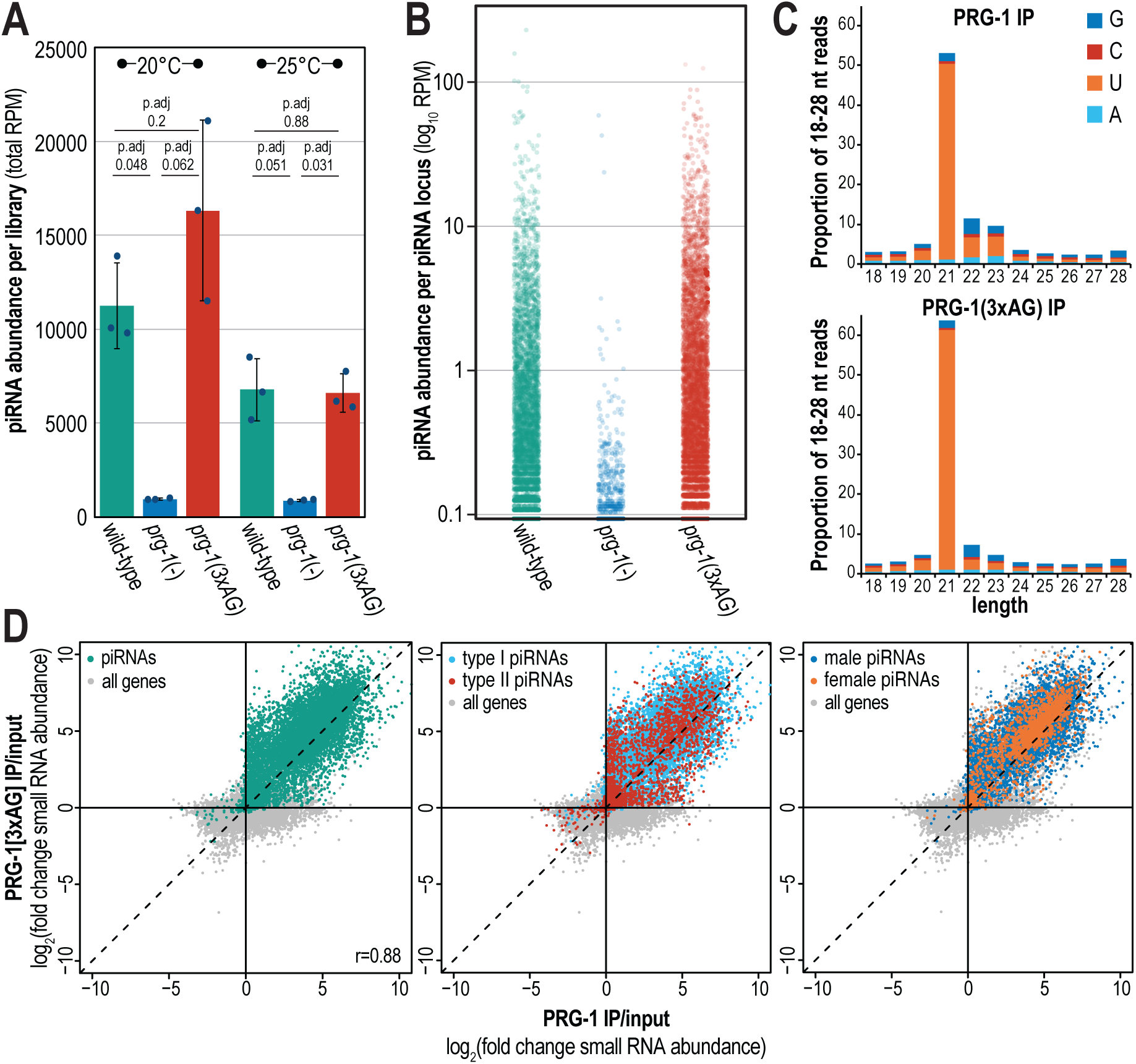
PRG-1 RG motifs are not required for piRNA production or loading. **A.** Total reads per million reads mapping to all piRNA gene loci in wild-type, *prg-1(null)* and *prg-1(3xAG)* mutants raised at either 20°C or 25°C indicate that piRNAs are not reduced in *prg-1(3xAG)* mutants. Error bars indicate standard deviation of three replicate libraries. Two-tail *t*-tests were performed to determine statistical significance and *p* values were adjusted for multiple comparisons. **B.** Reads per million reads mapping to each annotated piRNA in wild-type, *prg-1(null)* and *prg-1(3xAG)* mutants, where each dot is an individual piRNA gene locus. **C.** 5’ length and nucleotide distribution in mKate2::3xMyc::PRG-1 and in mKate2::3xMyc::PRG-1[3xAG] input and IP libraries, averaged from 3 replicate libraries. **D.** Scatterplot displaying log_2_(fold change) of small RNAs mapping to each gene in PRG-1 IP and PRG-1[3xAG] relative to input. piRNAs are teal (left panel), type I piRNAs are light blue (middle panel), type II piRNAs are red (middle panel), male piRNA are dark blue (right panel), female piRNAs are orange (right panel), and all other genes are gray (all panels).

Next, to further investigate the requirement for PRG-1 RG motifs, we sought to examine the impact of the RG motif mutant on the complement of small RNAs bound by PRG-1. To this end, we immunoprecipitated mKate2::3xMyc::PRG-1 and mKate2::3xMyc::PRG-1[3xAG] and generated high-throughput sequencing libraries of associated small RNAs. Both PRG-1 and PRG-1[3xAG] preferentially associate with 21-nt small RNAs with 5’ uridines, hallmarks of *C. elegans* piRNAs (Fig. 4C). Several proteins have been identified that are important for piRNA processing, including the exonuclease PARN-1 (Tang et al., 2016). In the *parn-1* mutant, piRNAs are not trimmed at their 3’ ends, resulting in an increased length distribution. No defects were observed in maturation or trimming of piRNAs in the PRG-1[3xAG] strain. Finally, to directly address if PRG-1 and PRG-1[3xAG] associate with the same piRNAs at similar levels, we compared the enrichment in each IP compared to input for small RNAs targeting each genic feature. We found that piRNAs were highly enriched in both PRG-1 and PRG-1[3xAG] IPs and the RNAs enriched in the two strains show a strong positive correlation (r=0.88) (Fig. 4D). Together, these data indicate that the PRG-1 RG motifs are not required for piRNA processing or PRG-1 loading.

piRNAs can be categorized based on the pathway from which they are generated and their sex-specific expression. Specifically, canonical (or type I) piRNAs, the majority of which reside in clusters on chromosome IV, have a conserved upstream “Ruby” motif and depend on PRDE-1 and SNPC-4 for expression; in contrast, motif-independent (type II) piRNAs are unclustered, lack the Ruby motif, and are generated independently of PRDE-1 and the SNPC complex (Gu et al., 2012; Kasper et al., 2014; Ruby et al., 2006; Weick et al., 2014). To determine whether disruption of the PRG-1 RG motifs are affects production, loading, or stability of type I vs type II piRNAs differently, we examined the enrichment of each type of piRNA in the wild-type PRG-1 and PRG-1[3xAG] IPs. We detected no differential enrichment of type I or type II piRNAs (Fig. 4D). Further, nucleotide differences in the Ruby motif and male-or female-specific SNPC proteins can promote the expression of male- and female-specific piRNAs (Billi et al., 2013; Choi et al., 2021). We similarly were unable to detect any difference in enrichment for male vs female piRNAs in the PRG-1 and PRG-1[3xAG] IPs. In summary, PRG-1 RG motifs are not required for production and processing of piRNAs, loading of piRNAs into PRG-1, or differential expression or loading of type 1/2 or male/female piRNAs, indicating that our observed phenotypic defects in the *prg-1(3xAG)* mutants must occur downstream of piRNA production and loading.

### PRG-1 RG motifs are required for production of downstream 22G-RNAs

piRNAs mediate target gene repression through the recruitment of the mutator complex and RdRPs, leading to the production 22G-RNAs (Bagijn et al., 2012; Lee et al., 2012). Since the RG motifs of PRG-1 are not required for piRNA production or loading, we next sought to investigate whether they are instead necessary for production of the mutator-dependent 22G-RNAs. To do so, we sequenced total small RNAs from both *prg-1(null)* and *prg-1(3xAG)* mutant adults, mapping them to the *C. elegans* genome. We observed significant changes in small RNA abundance for many genes in both *prg-1* mutants. Specifically, in the *prg-1(null)* mutant, small RNAs mapping to 654 genes significantly increased, while small RNAs mapping to 1781 genes significantly decreased (Fig. 5A). In the *prg-1(3xAG)* mutant, small RNAs mapping to 318 gene significantly increased, while small RNAs mapping to 478 genes significantly decreased (Fig. 5A). These data show that while piRNA levels were unaffected by the loss of PRG-1 RG motifs, other small RNAs were dramatically altered in abundance, with the effect being more pronounced in the *prg-1(null)* mutant compared to the *prg-1(3xAG)* mutant.

**Figure 5.**
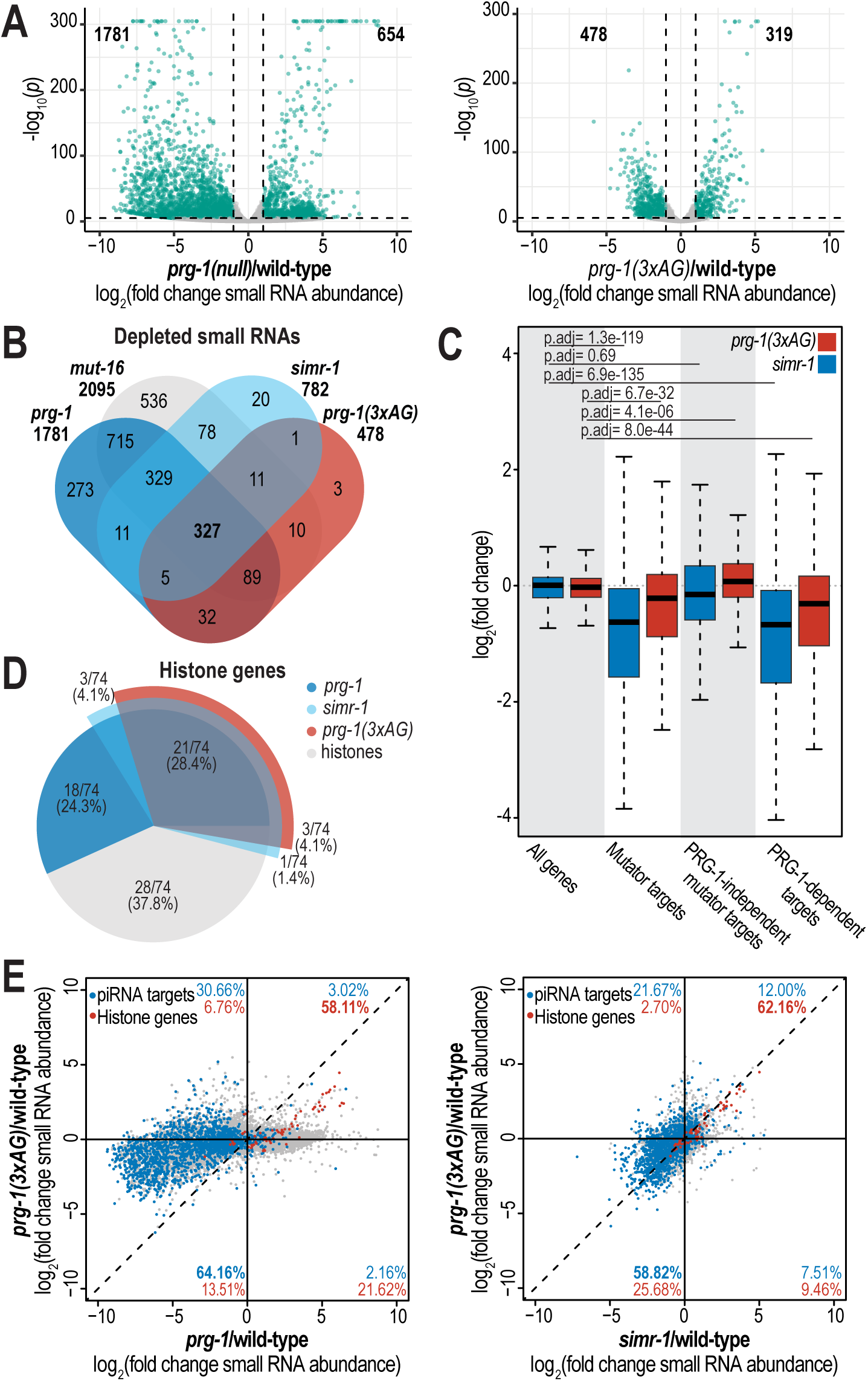
PRG-1 RG motifs are essential to carry out piRNA-dependent gene regulation. **A.** Volcano plot showing the log_2_(fold change) of small RNAs mapping to each *C. elegans* gene in *prg-1(null)* and *prg-1(3xAG)* mutants compared to wild-type animals (x-axis) and log_10_(adjusted *p* value) (y-axis). Genes with twofold change and adjusted *p* value of ≤0.05 are indicated in teal. Number of genes with significantly up- and down-regulated mapped small RNAs is indicated. **B.** Box plots displaying total small RNAs levels mapping to genes from the indicated small RNA pathways in *prg-1(3xAG)* and *simr-1* mutants compared to wild-type animals raised at 20°C. Details regarding definition of small RNA target gene classes is provided in the Materials and Methods section. Two-tail *t*-tests were performed to determine statistical significance and *p* values were adjusted for multiple comparisons. **C.** Venn diagrams indicating overlap of genes depleted of total small RNAs by two-fold or more in indicated mutants compared to wild-type. **D.** A pie chart of all histone genes shows the number of genes enriched for small RNAs in *prg-1*, *prg-1(3xAG)*, and *simr-1* mutants compared to wild-type. **E.** Scatterplot displaying log2(fold change) of small RNAs mapping to each gene in *prg-1(null)* or *simr-1* (x-axis) relative to *prg-1(3xAG)* (y-axis). PRG-1-dependent target genes (piRNA targets) are indicated in blue, histone genes are indicated in red, and all other genes are gray. Percentage of piRNA targets and histone genes in each quadrant are indicated.

The majority of mutator-target genes are also targeted by piRNAs (Manage et al., 2020). Therefore, we next examined whether the small RNAs depleted in the prg-1(3xAG) mutant overlap with those depleted in *prg-1(null)* and *mut-16* mutants, the latter of which completely abolishes the production of mutator-dependent 22G-RNAs. We found that 474 of 478 (99.2%) genes with reduced small RNAs in *prg-1(3xAG)* also had reduced small RNAs in *prg-1(null)* and/or *mut-16* mutants (Fig. 5B). Additionally, 416 of these genes (87.0%) had significantly reduced small RNAs in *both prg-1(null)* and *mut-16* mutants (Fig. 5B). Although piRNA-target genes account for the majority of mutator-targets, some mutator-dependent 22G-RNAs are still produced independent of PRG-1 and the piRNA pathway. To assess whether piRNA-target genes are more severely depleted of small RNAs in the *prg-1(3xAG)* mutant compared to other mutator-target genes, we analyzed a list of mutator-target genes whose small RNAs are unchanged or increased in *prg-1* mutants (Manage et al., 2020). These PRG-1-independent mutator targets showed no reduction in small RNA abundance in the *prg-1(3xAG)* mutant compared to wild-type; in fact, they exhibited a modest increase in small RNA levels (Fig. 5C). The PRG-1-independent mutator-targets were also significantly less depleted of small RNAs than either all mutator-target genes or piRNA-target genes. Together, these findings indicates that the PRG-1 RG motifs are important for the production of endogenous small RNAs at PRG-1-dependent mutator*-*target genes.

This disruption in small RNA production at PRG-1-dependent mutator targets in the *prg-1(3xAG)* mutant closely resembles a phenotype we previously observed in the Tudor-domain containing protein SIMR-1 mutant (Manage et al., 2020). Like *prg-1(3xAG)*, *simr-1* mutants exhibit a mortal germline, a defects in the reestablishment of WAGO-class 22G-RNAs, and a preferentially loss small RNAs at the PRG-1-dependent targets compared to PRG-1-independent mutator targets, yet they do not affect piRNA production or stability (Fig. 5C) (Manage et al., 2020). We therefore compared the genes with depleted small RNAs in *prg-1(3xAG)* mutants to those in *simr-1* mutants and found that 344 of 478 (72.0%) genes with reduced small RNAs in *prg-1(3xAG)* were also depleted in *simr-1*, and 327 of 478 (68.4%) were depleted in *prg-1(null)*, *mut-16*, and *simr-1* (Fig. 5B). Thus *prg-1(3xAG)* and *simr-1* share multiple phenotypes associated with piRNA gene regulation and contribute to the silencing of piRNA target genes, yet neither directly affects piRNA production or loading.

Enrichment of small RNAs at certain histone genes is a hallmark of disruptions in the piRNA pathway, but not the mutator pathway (Barucci et al., 2020; Manage et al., 2020; Montgomery et al., 2021; Reed et al., 2020). Upon examining the 74 *C. elegans* histone genes, we found that 24 (32.4%) were enriched for small RNAs in the *prg-1(3xAG)* mutant (Fig. 5D). In comparison, 28 (37.8%) were enriched in *simr-1* mutants and 42 (56.8%) were enriched in *prg-1(null)* mutants. Notably, all histone genes enriched for small RNAs in the *prg-1(3xAG)* mutant were also enriched in the *simr-1* mutant, further highlighting the similarity between these two mutants.

To further investigate the relationship between *prg-1(3xAG)* and *prg-1(null)* or *simr-1*, we compared the log2fold change in small RNA abundance in each mutant relative to wild-type. Despite the milder phenotype of the *prg-1(3xAG)* mutant compared *to prg-1(null)*, the two mutants showed a fairly strong correlation 64.2% of PRG-1-dependent target genes with reduced small RNAs were shared between the two mutants, and 58.1% of histone genes exhibited an increase in small RNA abundance (Fig. 5E). Similarly, the *prg-1(3xAG)* and *simr-1* mutants were also well correlated, with 58.8% of PRG-1-dependent target genes with reduced small RNAs shared between the two mutants and 62.2% of histone genes showing an increase in mapped reads (Fig. 5E). Taken together, these data suggest that the prg-1(3xAG) mutant modulates small RNA levels at PRG-1-dependent targets and histone genes in a manner similar to prg-1(null) and simr-1 mutants. This positions the function of PRG-1 RG motifs downstream of piRNA production and PRG-1 loading, and potentially in collaboration with SIMR-1, to mediate effective 22G-RNA biogenesis at piRNA target genes.

In this study, we investigated the functional role of the RG motifs in the N-terminal region of *C. elegans* PRG-1, focusing on their potential contribution to piRNA-mediated gene silencing and germline maintenance. Our results demonstrate that the RG motifs are not essential for PRG-1 expression, stability, or localization to germ granules, nor are they required for piRNA production, processing, or loading. However, mutations in these motifs result in defects in transgenerational fertility, germline atrophy at elevated temperatures, and impaired piRNA pathway function. Specifically, the RG motifs are critical for the proper production of downstream 22G-RNAs, essential for maintaining fertility and silencing of piRNA-targeted genes. Notably, our findings suggest that the RG motifs facilitate interactions necessary for the effective recruitment of downstream silencing machinery, underscoring the complex role of post-translational modifications in piRNA-mediated gene regulation. These results highlight the importance of the RG motifs in the PRG-1 protein for maintaining the integrity of the piRNA pathway, with broader implications for understanding the molecular mechanisms that govern transgenerational inheritance and gene silencing.

## Material and methods

### Strains

*C. elegans* strains were maintained at 20°C on NGM plates seeded with OP50 *E. coli* according to standard conditions unless otherwise stated (Brenner, 1974). All strains used in this project are listed in Supplementary Table 1.

### CRISPR-mediated strain construction

For the generation of *mKate2::3xMyc::prg-1(3xAG)*, the injection mix included 0.25 μg/μl Cas9 protein (IDT), 100 ng/μl tracrRNA, 14 ng/μl *dpy-10(cn64)* crRNA, 42 ng/μl *prg-1* crRNA, and 110 ng/μl of the oligo repair template were injected into USC1232 (*prg-1(cmp220[(mKate2 + loxP + 3xMyc)::prg-1])*) (Dokshin et al., 2018; Paix et al., 2015). For the generation of *prg-1* mutants in the piRNA sensor strain, the injection mix included 0.25 μg/μl Cas9 protein (IDT), 100 ng/μl tracrRNA, 14 ng/μl *dpy-10(cn64)* crRNA, 42 ng/μl *prg-1* crRNA, and 110 ng/μl of the appropriate *prg-1* mutant oligo repair template were injected into HCL-134(*oma-1::gfp I; cdk-1::gfp II; 21ur-anti-gfp(type I) IV*) (Brown et al., 2023). Guide RNA sequences are provided in Supplementary Table 2.

### Immunoprecipitation and Mass Spectrometry

For immunoprecipitation experiments, ∼1 million synchronized mKate::3xMyc::PRG-1 (strain USC1232) adult animals (∼68 hrs at 20 °C after L1 arrest) were collected in IP Buffer (50 mM Tris-Cl pH 7.4, 100 mM KCl, 2.5 mM MgCl2, 0.1% Igapal CA-630, 0.5 mM PMSF, cOmplete Protease Inhibitor Cocktail (Roche 04693159001)) frozen in liquid nitrogen, and homogenized using a mortar and pestle. After further dilution into IP buffer (1:10 packed worms:buffer), insoluble particulate was removed by centrifugation and 10% of sample was taken as “input.” The remaining lysate was used for the immunoprecipitation. Immunoprecipitation was performed at 4 °C for 1 h with pre-conjugated anti-Myc affinity matrix (Thermo Fisher 20168), then washed at least three times in immunoprecipitation buffer. A fraction of each sample was analyzed by western blot to confirm efficacy of immunoprecipitation. 2x sample buffer was added to the remainder of each sample, followed by gel electrophoresis (4–12% Bis-Tris polyacrylamide gels, Thermo Fisher) and overnight colloidal Coomassie staining.

Bands containing immunoprecipitated protein were excised from gel and cut into ∼1 mm3 pieces. Gel pieces were then subjected to a modified in-gel chymotrypsin digestion procedure (Shevchenko et al., 1996). Gel pieces were washed and dehydrated with acetonitrile for 10 min. followed by removal of acetonitrile. Pieces were then completely dried in a speed-vac. Rehydration of the gel pieces was with 50 mM ammonium bicarbonate solution containing 12.5 ng/µl modified sequencing-grade chymotrypsin (Promega) at 4 °C. After 45 min, the excess chymotrypsin solution was removed and replaced with 50 mM ammonium bicarbonate solution to just cover the gel pieces. Samples were then incubated at room temperature overnight. Peptides were later extracted by removing the ammonium bicarbonate solution, followed by one wash with a solution containing 50% acetonitrile and 1% formic acid. The extracts were dried in a speed-vac (∼1 h) and stored at 4 °C until analysis.

On the day of analysis, the samples were reconstituted in 5–10 µl of HPLC solvent A (2.5% acetonitrile, 0.1% formic acid). A nano-scale reverse-phase HPLC capillary column was created by packing 2.6 µm C18 spherical silica beads into a fused silica capillary (100 µm inner diameter x ∼30 cm length) with a flame-drawn tip (Peng and Gygi, 2001). After equilibrating the column each sample was loaded via a Famos auto sampler (LC Packings) onto the column. A gradient was formed and peptides were eluted with increasing concentrations of solvent B (97.5% acetonitrile and 0.1% formic acid).

As peptides eluted, they were subjected to electrospray ionization and then entered into an LTQ Orbitrap Velos Pro ion-trap mass spectrometer (Thermo Fisher). Peptides were detected, isolated, and fragmented by collision-induced dissociation to produce a tandem mass spectrum of specific fragment ions for each peptide. Peptide sequences (and hence protein identity) were determined by matching protein databases with the acquired fragmentation pattern by the software program, Sequest (Thermo Fisher). All databases include a reversed version of all the sequences and the data was filtered to between a one and two percent peptide false discovery rate.

### Imaging and image quantification

All animals were imaged as adults, 68 hours post hatching, and immobilized in ddH2O with 0.5 M sodium azide before mounting on glass slides for imaging.

Colocalization analysis: Images were collected on the Lecia Stellaris 5 confocal microscope using the 63x plan apo N.A 1.40 oil immersion objective. Z stacks were obtained with a 0.2 μM step size. 5 Z slices were then maximum projected using FIJI. Quantitative colocalization analysis between different granules was performed in FIJI/ImageJ2 using the Coloc2 package. At least 3 individual nuclei from each gonad, and at least 10 individual gonads for a total of at least 30 nuclei were used to calculate Pearson’s R value.

Germline Length: Images were collected on the Deltavision Elite widefield microscope (GE Healthcare) using the 60x N.A 1.42 oil immersion objective. Z stacks were obtained with a 0.2 μM step size. Germline length was measured using the multisegmented line tool in FIJI. Measurements were made from the mitotic tip to the bend of the gonad.

piRNA sensor fluorescence intensity: Images were collected on the Deltavision Elite widefield microscope (GE Healthcare) using the 60x N.A 1.42 oil immersion objective. Z stacks were obtained with a 0.2 μM step size. 5 Z slices were then maximum projected using FIJI. A standardized region of interest (ROI) of 70x70 pixels was then used to measure the fluorescence intensity of the -1 oocyte nucleus and the background fluorescence. The following formula was used to determine the corrected total cellular fluorescence (CTCF) of the -1 oocyte nucleus:

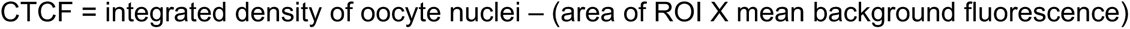

### Transgenerational fertility assay

Wild-type and mutant *C. elegans* strains were maintained at 20°C prior to transgenerational fertility assays. Animals were shifted to 25°C at L4 and allowed to lay (P0). A single progeny from each plate was selected and moved to a new plate at the L4 stage for the following generation. Lines considered to be sterile were kept for an additional 3 days to ensure no eggs would be laid or hatched. This was conducted until all lines of *prg-1(3xAG)* and *prg-1(n4357)* animals were sterile.

### Reestablishing WAGO-class 22G-siRNA production

The mutator pathway was restored to WAGO-class 22G-RNA-defective animals according to the crossing scheme in Fig. 3C and as previously described (Phillips et al., 2015). The *unc-119* mutation was always present in the parental hermaphrodite strain to allow for unambiguous identification of cross vs. self progeny. F1 animals were singled to individual plates as L4 larvae and scored 2–3 days later for presence or absence of progeny.

### Western blots

Synchronized adult *C. elegans* were harvested (∼68 hrs at 20 °C after L1 arrest) and 200 adults were loaded per lane. Proteins were resolved on 4–12% Bis-Tris polyacrylamide gels (ThermoFisher), transferred to nitrocellulose membranes, and probed with anti-actin 1:10,000 (Abcam ab3280), or anti-Myc 1:1,000 [9E10 clone] (ThermoFisher 13–2500). Secondary HRP antibodies were purchased from ThermoFisher.

### Small RNA library preparation

For immunoprecipitation followed by small RNA sequencing, ∼100,000 synchronized adult animals (∼68 h at 20 °C after L1 arrest) were washed off plates with H2O and collected in IP Buffer (50 mM Tris-Cl pH 7.5, 100 mM KCl, 2.5 mM MgCl2, 0.1% Nonidet P40 substitute) containing Protease Inhibitor (Thermo Fisher A32965) and RNaseOUT Ribonuclease Inhibitor (Thermo Fisher 10777019). Immunoprecipitation was performed as described above using anti-Myc Affinity Matrix (ThermoFischer, 20168). PRG-1 or PRG-1[3xAG]-bound RNAs were isolated using TRIzol reagent (Thermo Fisher, 15596018), followed by chloroform extraction and isopropanol precipitation. Small RNAs (18 to 30-nt) were size selected on denaturing 10% polyacrylamide gels from total RNA samples. Small RNAs were treated with 5′ RNA polyphosphatase (Epicentere RP8092H) (total small RNA samples) or 5’ RNA pyrophosphohydrolase (NEB M0356S) (IP samples) and ligated to 3′ pre-adenylated adapters with Truncated T4 RNA ligase (NEB M0373L). Small RNAs were then hybridized to the reverse transcription primer, ligated to the 5′ adapter with T4 RNA ligase (NEB M0204L), and reverse transcribed with Superscript III (Thermo Fisher 18080-051). Small RNA libraries were amplified using Q5 High-Fidelity DNA polymerase (NEB M0491L) and size selected on a 10% polyacrylamide gel. Library concentration was determined using the Qubit 1X dsDNA HS Assay kit (Thermo Fisher Q33231) and quality was assessed using the Agilent BioAnalyzer. Libraries were sequenced on the Illumina NextSeq500 (SE 75-bp reads) platform. Primer sequences are available in Supplementary Table 2. Differentially expressed gene lists can be found in Supplementary Table 3. Sequencing library statistics summary can be found in Supplementary Table 4.

### Bioinformatic analysis

For small RNA libraries, sequences were parsed from adapters using FASTQ/A Clipper (options: -Q33 -l 17 -c -n -a TGGAATTCTCGGGTGCCAAGG) and quality filtered using the FASTQ Quality Filter (options: -Q33 -q 27 -p 65) from the FASTX-Toolkit (version 0.0.13) (http://hannonlab.cshl.edu/fastx_toolkit/), mapped to the *C. elegans* genome WS258 using Bowtie2 v. 2.2.2 (default parameters) (Langmead and Salzberg, 2012), and reads were assigned to genomic features using FeatureCounts (options: -t exon -g gene_id -O --fraction – largestOverlap) which is part of the Subread package (version 2.0.1) (Liao et al., 2014, 2013). Differential expression analysis was performed using DESeq2 (version 1.22.2) (Love et al., 2014). To define genes depleted of small RNAs in *prg-1(null)* and *prg-1(3xAG)* mutants, a twofold-change cutoff, a DESeq2 adjusted *p* value of ≤0.05, and at least 10 RPM in the wild-type or mutant libraries were required to identify genes with significant changes in small RNA levels.

Type II piRNAs, male and female piRNAs, mutator targets, PRG-1-dependent targets, PRG-1-independent mutator targets, and histone genes were previously described (Gu et al., 2012; Choi et al., 2021; Manage et al., 2020; Pettitt et al., 2002). Additional data analysis was done using R, Excel, and Python and modified in Adobe Illustrator. Sequencing data is summarized in Supplementary Table 3.

## Data Availability

The RNA sequencing data generated in this study are available through Gene Expression Omnibus (GEO) under accession code GSE138220 for *simr-1* data and GSE296607 for all other data.

## Acknowledgements

We thank the members of the Phillips lab for helpful discussions and feedback on the manuscript and the lab Heng-Chi Lee for generously providing strains. This work was supported by the National Institute of Health grant R35 GM119656 (to CMP) and Dylan Wallis has been supported as a USC Dornsife-funded Chemistry-Biology Interface trainee. Some strains were provided by the CGC, which is funded by NIH Office of Research Infrastructure Programs (P40 OD010440). Next generation sequencing was performed by the USC Molecular Genomics Core, which is supported by award number P30 CA014089 from the National Cancer Institute.

## Author contributions

D.W..: Conceptualization, Investigation, Formal analysis, Writing–original draft, Writing–reviewing and editing, Visualization

C.M.P.: Conceptualization, Formal Analysis, Writing–original draft, Writing–reviewing and editing, Supervision, Funding Acquisition.

---

**Figure S1.**
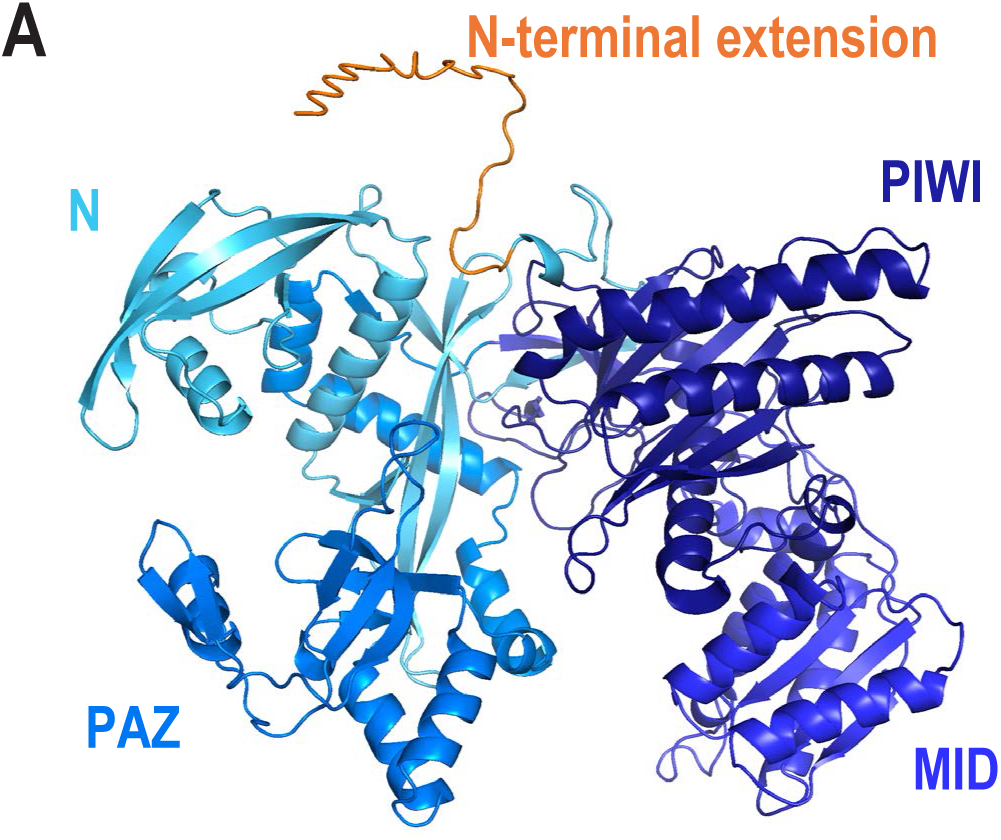
Predicted Structure of PRG-1. A. Alphafold2 predicted structure of PRG-1 with the N-terminal extension shown in orange. Structured domains of PRG-1 are indicated in various shades of blue.

